# RNA Polymerase II is involved in 18S and 25S ribosomal RNA transcription, in *Candida albicans*

**DOI:** 10.1101/510156

**Authors:** Jacob Fleischmann, Miguel A. Rocha, Peter V Hauser

**Affiliations:** Research Division, Greater Los Angeles VA Healthcare System, Los Angeles, California, USA; Department of Medicine, David Geffen School of Medicine at UCLA, Los Angeles, California, USA.; Department of Integrative Biology and Physiology, University of California at Los Angeles, Los Angeles, California, USA.

## Introduction

While in eukaryotes three RNA polymerases are involved in ribosome production [1] under usual growth conditions, the 18S and 25S ribosomal RNA (rRNA) components are thought to be exclusively the products of transcription by RNA polymerase I (Pol I) followed by processing [2]. We have observed recently, in *Candida albicans* during nutritional depletion and with TOR inhibition, the appearance of 18S and 25S rRNA molecules, resisting digestion by a 5′-phosphate-dependent exonuclease, indicating that they were different from the usual processed rRNA transcripts [3]. *Candida albicans*, a eukaryotic yeast, is a major cause of invasive fungal disease especially in immune compromised patients [4]. Ribosomes of eukaryotic cells are assembled from four individual rRNAs and 79 proteins [5]. As in *Saccharomyces cerevisiae*, genes coding for rRNA (rDNA) in *C. albicans* are repeated multiple times in tandem [6], allowing for efficient transcription by Pol I. Like other eukaryotes, the current accepted mechanism of the production of the 18S and 25S components of the ribosome in this yeast, is transcription of a 35S copy of the rDNA, followed by post and co-transcriptional processing of the nascent RNA [7]. Typically, processed RNA molecules will have a single phosphate on their 5’-end making them vulnerable to processive 5′→3′ exonucleases (P53E) that digests only RNA that has a 5′-monophosphate end [8]. Therefore, after digestion by such an exonuclease, it was unexpected to find 18S and 25S rRNA molecules in total RNA isolated from *C. albicans* entering its stationary phase [3]. Similar molecules were appearing also in yeast, whose TOR was inhibited by rapamycin [3]. This background information is illustrated in Fig 1A. Pyrophosphatase digestion which separates linked phosphates, made these resistant 18S and 25S molecules vulnerable again to 5’-exonulease digestion [3]. This indicated that these molecules contained more than a single phosphate at their 5’-end. This in turn raised the possibility that they were newly transcribed rather than processed, as polymerases use triphosphate nucleotides when they initiate transcription. Another serial enzyme digestion included alkaline phosphatase (AP) followed by P53E digestion, in this case the rRNAs remained protected [3]. This could have been due to either more than one phosphate being digested by AP resulting in 5′-OH, or further modification in rRNA, such as a 5’-cap protecting against both AP and P53E, again preventing exonuclease digestion. Additionally, we have previously found that *C. albicans* grown overnight, polyadenylates some of its 18S [9] and 25S [10] rRNA molecules, a feature associated with Pol II transcription [11]. These features prompted us to see whether Pol II is involved with ribosomal rRNA transcription.

**Fig 1.**
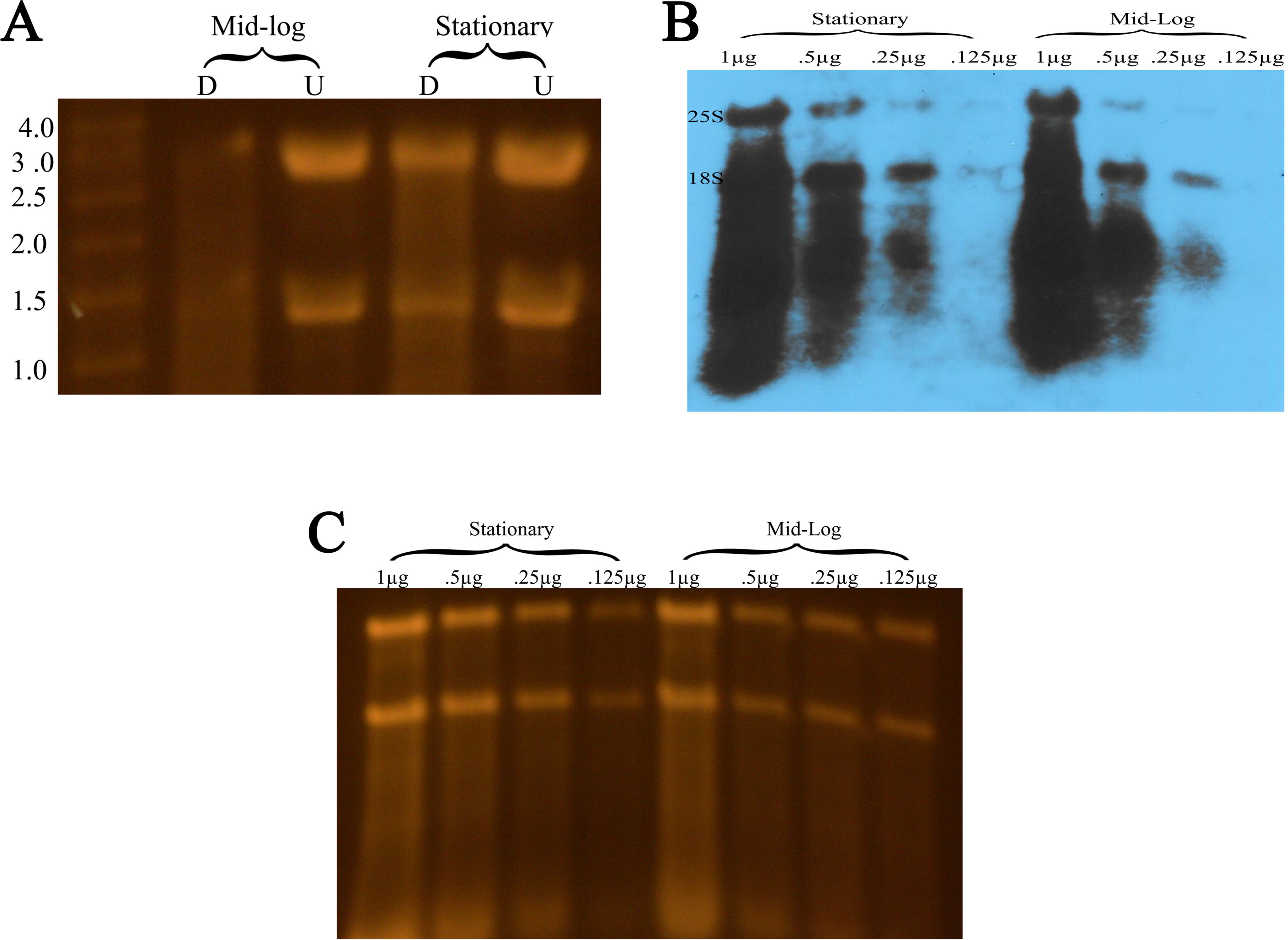
Ribosomal RNA molecules resistant to P53E in stationary phase C. *albicans* analyzed with anti-m7G-cap mAb. (A) Representative SYBR gold stained agarose gel depicting total RNA extracted from mid-log and stationary C. *albicans* that was digested (D) by P53E or undigested (U) showing significant percentage of both 18S and 25S being resistant to P53E. (B) Immunoblot of serial diluted total RNA (stationary and mid-log) using anti-m7G-cap mAb. This antibody has the strongest affinity for 7-methylguanosine cap but will also react with m7G within RNA (S1 Fig) (https://www.mblintl.com/products/rn016m). RNA from stationary cells show more intense signals for each dilution and continue to show signal at a dilution where RNA from mid-log no longer shows any signal. (C) Agarose gel from which immunoblotting was done, showing that the differences of immunoblot band intensities were not related to differences in RNA loading.

## Results

All transcripts synthesized by Pol II are distinguished by the presence of a 5’cap [12]. No such capping has been described for Pol I products. Thus, our approach was to see if any 18S and 25S transcripts produced by this yeast contained a 5’cap. We first utilized a monoclonal anti-7-methylguanosine (m^7^G) antibody (m^7^G-AB). This antibody is capable of cross-reacting with m^7^G within RNA, but has the highest affinity for 7-methylguanosine diphosphate (m^7^Gpp) attached to RNA by a 5’-5’triphosphate linkage (S1 Fig). To take advantage of this extra affinity, we extracted total RNA from yeast in mid-log growth phase (5-6 hours) and from those approaching stationary phase (12-16 hours). Same amount of RNA was taken from each time point and was serially diluted prior to gel electrophoresis and immunoblotting. As can be seen Fig 1B, m^7^G-AB detected more molecules for both 18S and 25S in each dilution from stationary phase RNA. Furthermore, it detected molecules in more dilutions from stationary RNA. Accompanying gel (Fig 1C) shows that these differences were not related to gel loading. We have persistently observed in multiple immunoblots, stronger signals for 18S than 25S, which suggests that these molecules were indeed not produced in equal amounts in a polycistronic fashion. The specificity of our antibody for 5’cap was also indicated by dot-blot analysis, as we could block the antibody with purchased m^7^Gpp (S2 Fig).

RNA immune precipitation (RIP) was also performed with the same antibody used in immunoblotting. Results of q-PCR amplifications of reverse transcribed RNAs that were obtained from the precipitations, are seen in Fig 2. While there was some amplification of 18S and 25S in RNA from mid-log phase organisms, likely from antibody cross-reacting within the RNA, amplification in RNA from stationary phase was more robust due to the additional affinity for the 5’cap.

**Fig 2.**
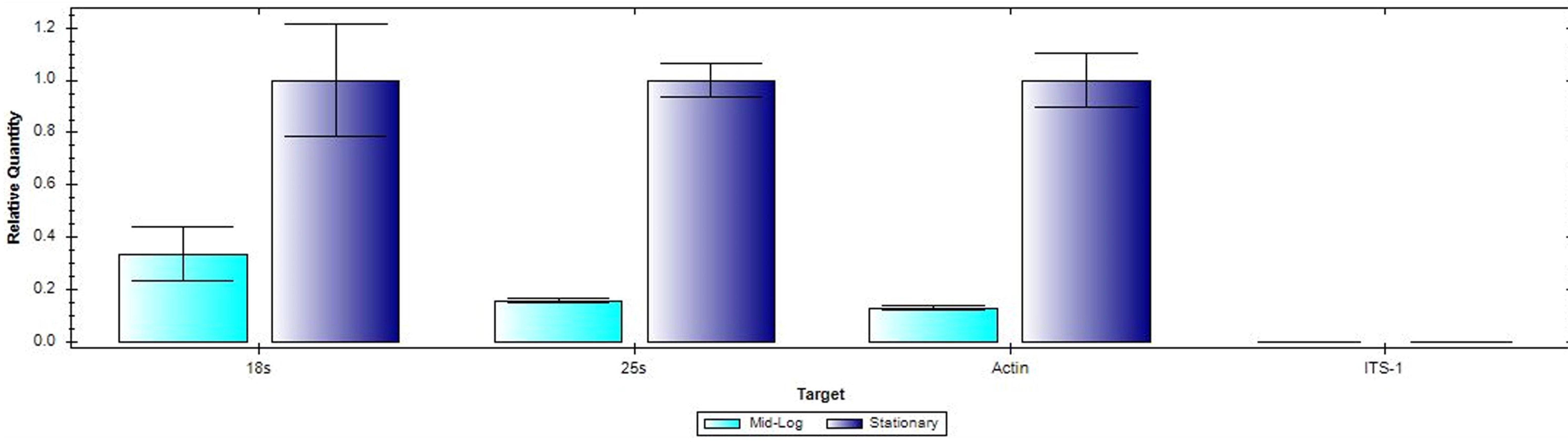
RIP and qPCR analysis. RT-qPCR quantification of 18S and 25S molecules precipitated from total RNA by anti-m7G-cap mAb, both from mid-log and stationary organisms. Positive and negative controls were actin and ITS-1 specific primers respectively. Error bars represent standard deviation (+/−) from three different experiments.

His-tagged EIF4E, a subunit of eIF4F cap binding protein complex (CBP) [13] was also used to precipitate any capped 18S and 25S molecules. EIF4F has special affinity for 5’cap and is unlikely to be cross reacting with parts of an RNA molecule other than the 5’cap. RNA from stationary cells indeed contained such molecules and none could be seen in RNA from mid-log organisms (Fig 3A). Anti-cap antibody reacted to these molecules (Fig 3B), further confirming that they contained 5’cap. Removing the cap by a pyrophosphatase reaction prevented their precipitation (Fig 3C), again substantiating the presence of a 5’cap on them. q-PCR amplification was performed on these precipitated molecules. The specificity of the cap binding protein is reinforced as amplification occurred primarily from molecules obtained from stationary organism (Fig 4) and very little from mid-log organisms.

**Fig 3.**
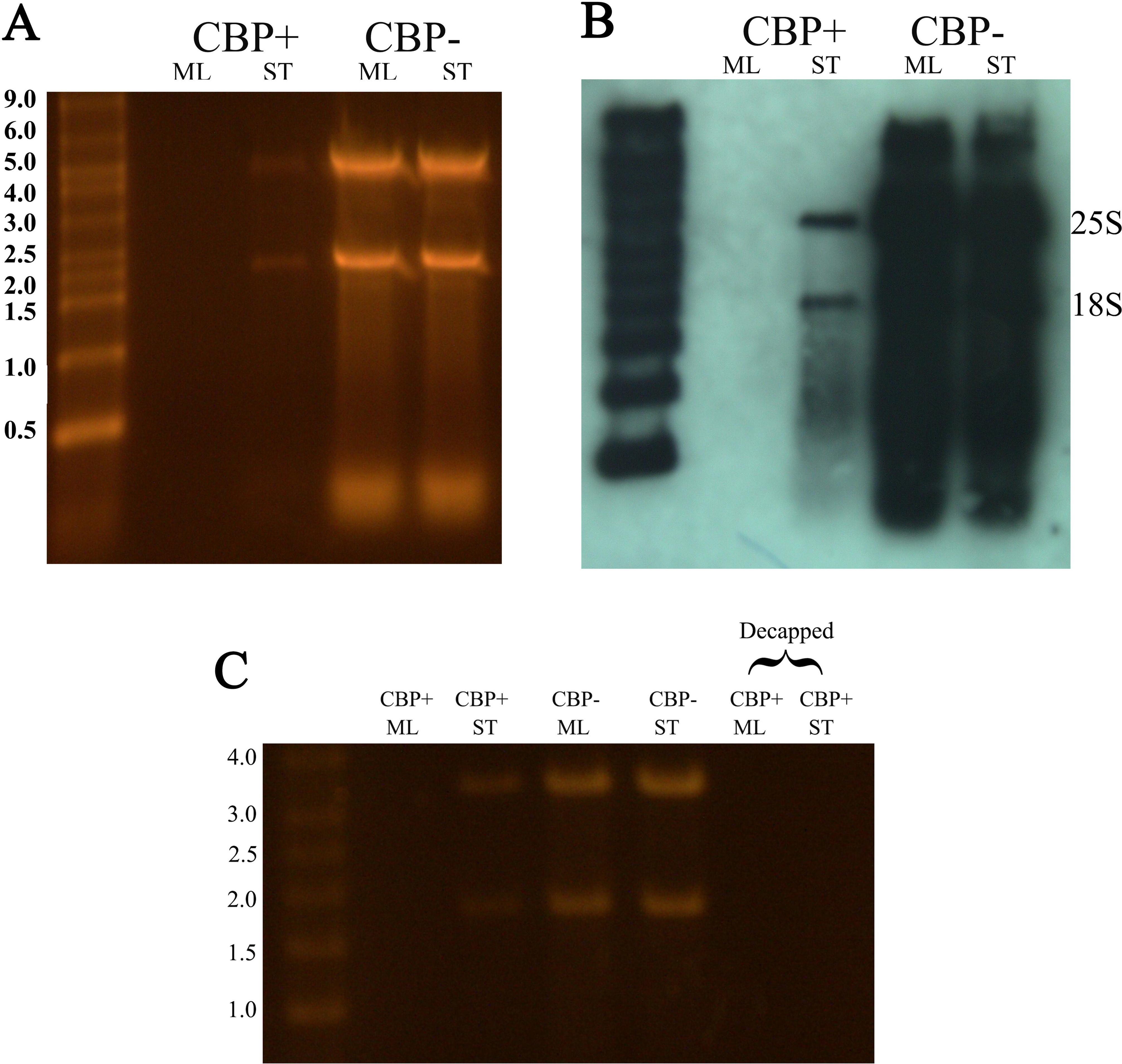
Ribosomal RNA molecules resistant to P53E in stationary phase C. *albicans* analyzed with cap binding protein eIF4e. (A) SYBR gold stained gel showing total RNA extracted from mid-log (ML) and stationary (ST) C. *albicans*. CBP-indicates total RNA without CBP precipitation and CBP+ represents RNA precipitated by CBP. (B) Immunoblot using anti-m7G-cap mAb to detect bands precipitated by CBP. The affinity for bands seen in CBP+ ST is confirming that it is a 5’-cap. Such capped molecules are detected only in RNA obtained from stationary yeast and none in RNA from mid-log organisms. The intensity of signal in CBP- is related to cross-reaction with m7G within RNA (https://www.mblintl.com/products/rn016m). (C) In addition to the conditions in Fig 2A, decapping prior to CBP precipitation, eliminates CBP from pulling out bands seen in CBP+ ST containing the cap.

**Fig 4.**
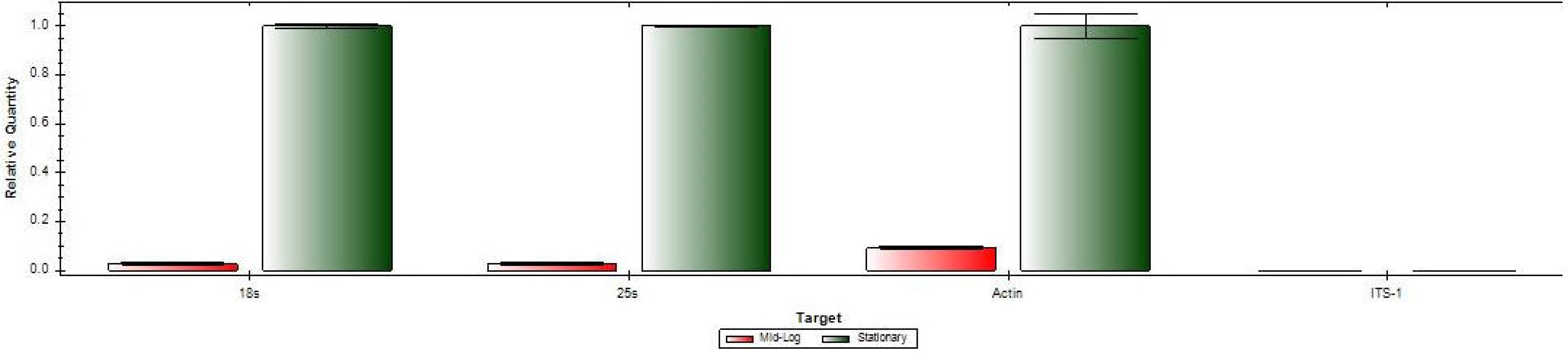
Cap binding protein precipitation and qPCR. RT-qPCR quantification of 18S and 25S molecules precipitated from total RNA by CBP both from mid-log and stationary organisms. Positive and negative controls were actin and ITS-1 specific primers respectively. Error bars represent standard deviation (+/−) from three different experiments.

As Northern blot analysis show these exonuclease resistant 18S and 25S molecules to be the same size as processed ones we carried out oligo ligation and amplification to confirm where the start sites of these molecules are. As can be seen in Fig 5A we were able to amplify appropriate PCR products with primers specific for oligo and 18S and 25S sequences even after combined CIP and Cap-Clip™ (similar in function to tobacco acid pyrophosphatase TAP) digestion. CIP digestion alone prevented any such amplification, indicating that the enzyme eliminated the 5’ phosphate completely. Thus, CAP-Clip™ treatment following CIP digestion allowed the gamma phosphate of cap to be exposed for ligation. Sequencing showed to be exactly at the processing site both for 18S and 25S (Fig 5B).

**Fig 5.**
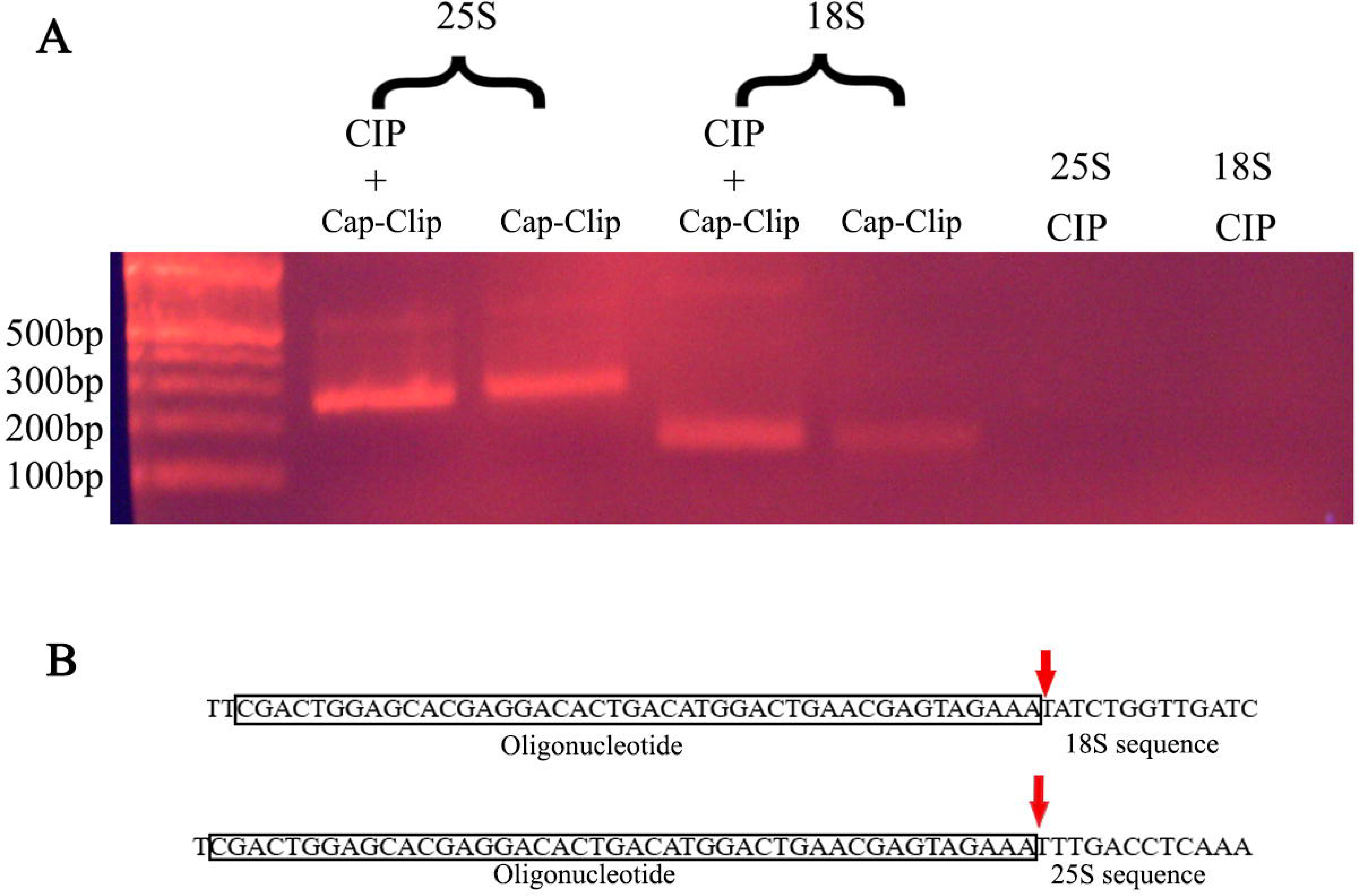
5’ end analysis of exonuclease resistant 18S and 25S. (A) Stained gel of PCR products amplified with oligo and internal specific primers (see ST table 1) for 18S and 25S. No amplification was obtained for CIP alone samples. (B) Sequences of 18S and 25S from CIP + CAP-Clip™ amplicons. Red arrow indicates start of processing site.

To see if capped 18S and 25S are also polyadenylated, we did poly-A selection on total RNA from both mid-log and stationary organism followed by immunoblotting with anti-cap antibody. As can be seen in Fig 6A the SYBR gold stained gel shows the presence of both 18S and 25S after poly-A selection from stationary organisms but not from mid-log organisms. Immunoblotting confirms 5’-cap on both 18S and 25S, though 18S is more difficult to see due to interference from mRNAs similar in size being detected by the antibody (Fig 6B). We also confirmed polyadenylation from the opposite direction. RNA precipitated with cap binding protein was reverse transcribed with a poly-T primer and amplified with appropriate primers. As can be seen in Fig 6C there are cap-selected molecules polyadenylated. The polyadenylation site for 18S was downstream from the reported processing site, but for 25S it was at the processing site. This is similar to what we found previously with 18S [10] and 25S [9] molecules from *C. albicans.*

**Fig 6.**
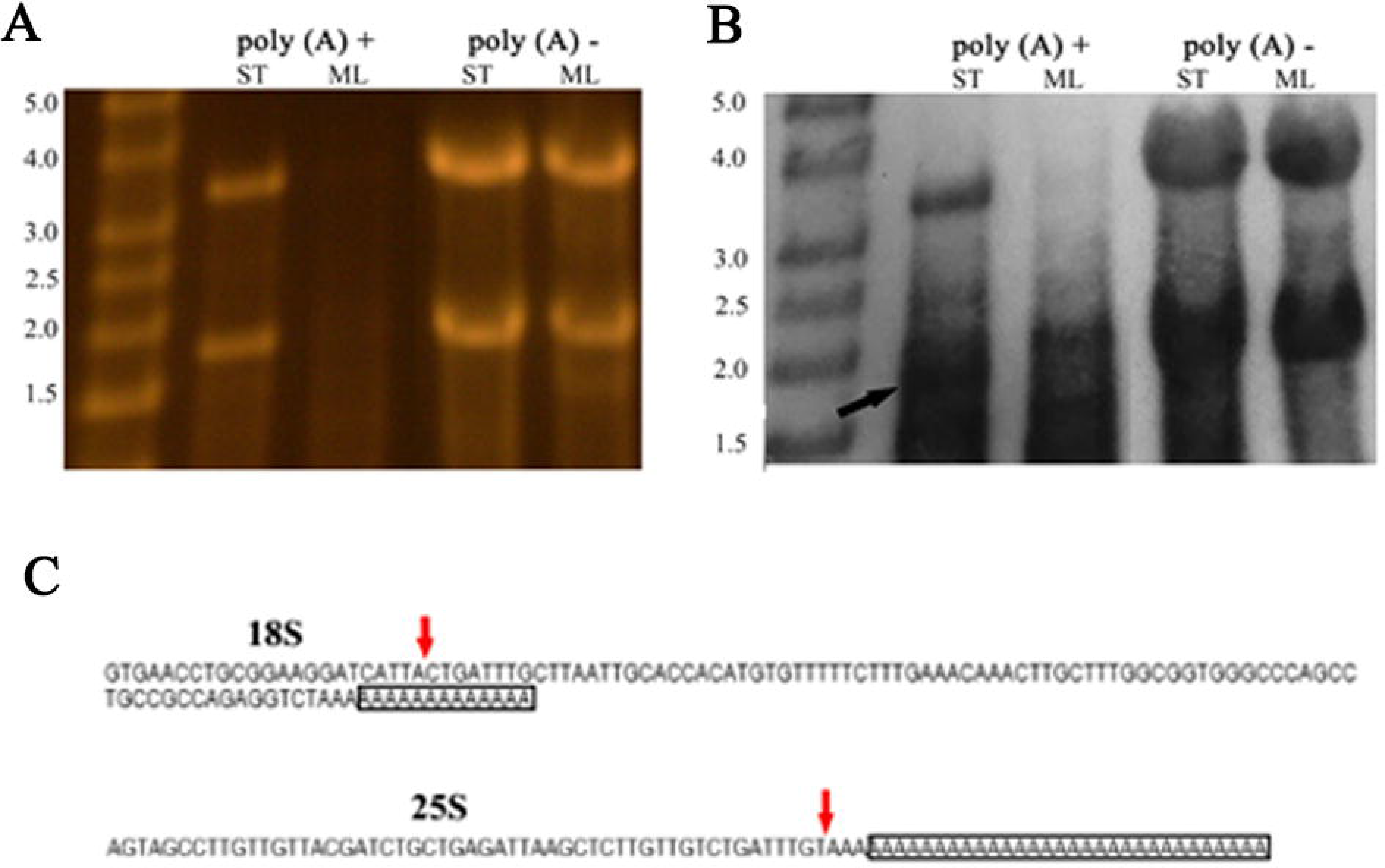
18S and 25S ribosomal RNA molecules with caps and polyadenylation isolated from stationary C. albicans. (A) SYBR gold stained gel with total RNA from mid-log (ML) and stationary (ST) C. *albicans*, before and after poly A selection using oligo dT beads showing significant polyadenylation of both 18S and 25S rRNAs. (B) immunoblot using anti-m7G-cap mAb to detect capped RNA before and after poly A selection. The same gel shown in Fig 3A was used for the immunoblotting. The arrow indicates the position of 18S seen in Fig 3A. (C) Sequences of 18S and 25S RNA from stationary yeast reverse transcribed with a poly dT primer (PI and PJ, see S1 Table. The RNA used was precipitated with CBP. Red arrows indicate reported processing site. Boxed letters indicate the length of the poly-T primers.

To show involvement by Pol II in rRNA transcription, we utilized chromatin immune precipitation (ChIP) and direct nucleolar visualization by fluorescence microscopy. Total DNA isolated from stationary *C. albicans* was precipitated with an antibody that recognizes the largest subunit of RNA polymerase II (Pol II-AB) [14] and the precipitated DNA was amplified with 18S and 25S specific primers (Fig 7). PCR products of predicted sizes were obtained from DNA isolated from stationary organisms and were sequenced to confirm that they were indeed 18S and 25S molecules. The same primers did not amplify any PCR products from DNA obtained from mid-log organisms.

**Fig 7.**
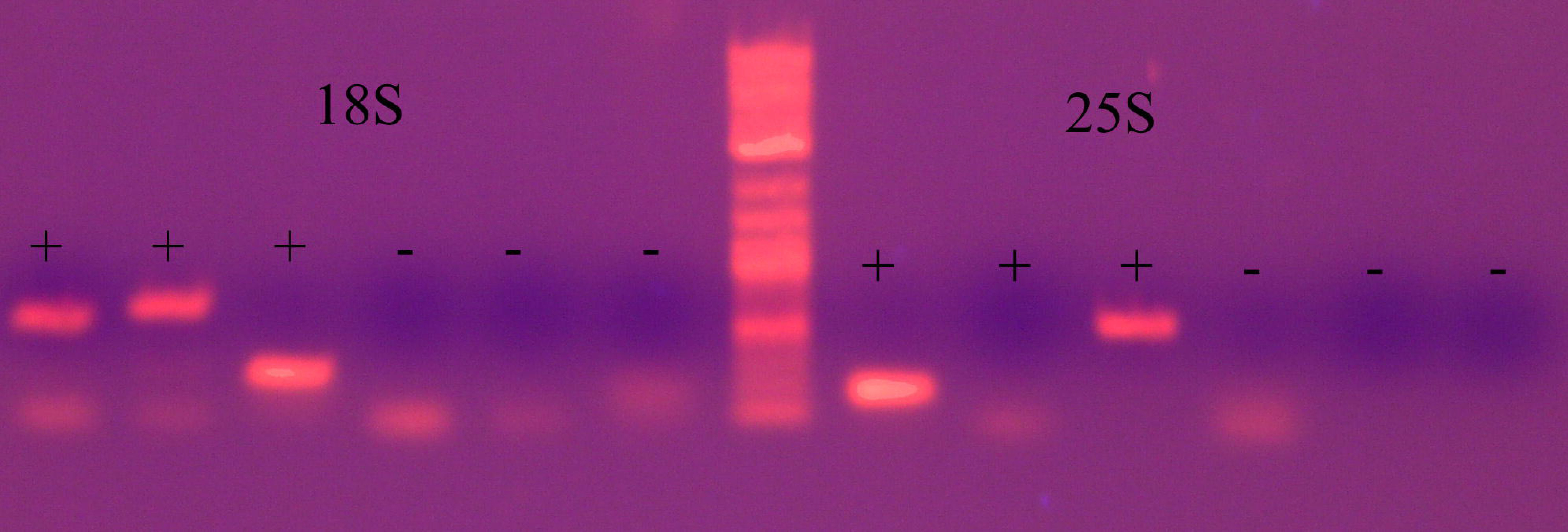
Evidence for the role of RNA Pol II in the transcription of 18S and 25S molecules in stationary C. *albicans*. (A) Chromatin Immunoprecipitation (ChIP) with polymerase II specific antibody. PCR fragments amplified from stationary organisms. Cells were cross-linked and chromatin was sheared by sonification. RNA Polymerase II mAb CTD4H8 (Epigentek) was used to precipitate DNA-protein complex. PCR was performed using three different sets of specific primers for 18S (PO-PB, PA-PP, PK-PQ) and 25S (PR-PD, PC-PS, PL-PT) (+). See Supplementary Table 1 for primers information. A non-immune IgG antibody was used as negative control (−).

Nucleoli of eukaryotic cells are the visible parts of nuclei where ribosomal DNA repeats are concentrated [15] and is thought to be the exclusive domain of Pol I [16]. Careful zymolase digestion of yeast cell walls [17] allowed us to use antibodies to pinpoint Pol II in nucleoli. As can be seen in Fig 8D, cells in stationary phase, many but not all, whose nucleoli are pinpointed by anti-Nop1p antibody [18] also stain over the same spots with Pol II-AB, detecting their presence and indicating that these cells are utilizing Pol II in ribosomal RNA production.

**Fig 8.**
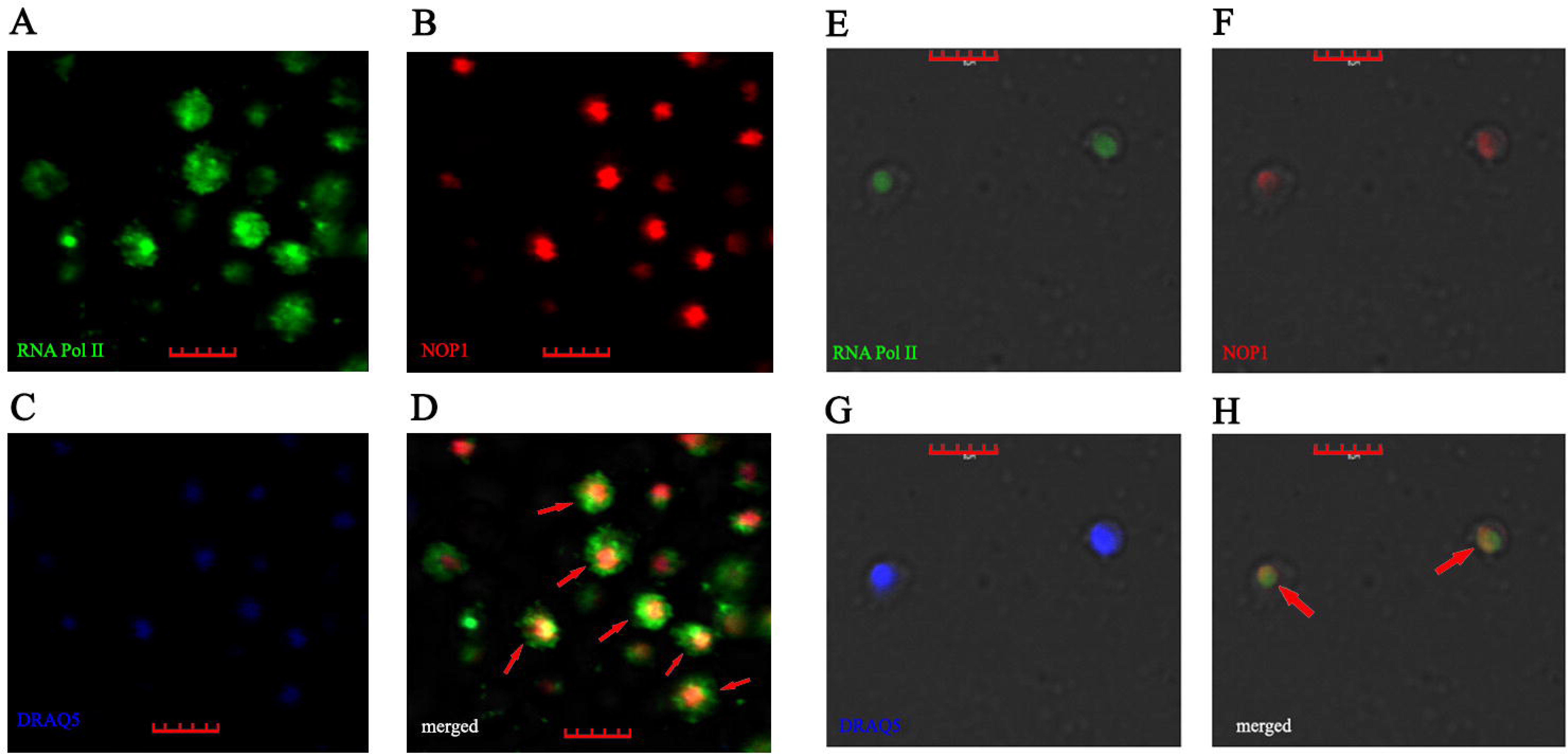
Immunofluorescence microscopy of nucleoli. (A-D) Micrographs of stationary C. *albicans* showing the presence of Pol II in the nucleolus. (B) primary antibody was specific for the c-terminal component of RNA PolII and secondary antibody was anti-mouse-alexa-488. (B) primary antibody was specific for NOP1 and the secondary was anti-mouse-alexa-568. (C) Nuclei stained with DRAQ5. (D) panels (A) and (B) merged. Red arrows point to nucleoli with the presence of Pol II. (E-H) Micrographs of rapamycin-inhibited C. *albicans.* The staining process for the micrographs is the same as in (A-D). Scale bars 5μm

## Discussion

Our data show that *C. albicans* produces 18S and 25S rRNA molecules with 5’-caps and poly-A tails, two features associated with Pol II. This combined with ChIP analysis and fluorescence microscopy showing the presence of Pol II at rRNA repeats, establishes that they were indeed newly copied by this enzyme. The 5’-cap explains why we found molecules that were resistant to 5′-phosphate-dependent exonuclease digestion^3^. While ribosome generation has not been a focus in *C. albicans*, studies in *Saccharomyces cerevisiae* and other yeast has been extensive and should be relevant to our organism. That Pol II is capable of transcribing rRNA has been shown in *S. cerevisiae.* When *RRN9*, one component of the Pol I upstream activating factor (UAF) was deleted inactivating Pol I, the yeast was capable of ribosome production utilizing Pol II, initiating transcription upstream from the normal Pol I start site still in a polycistronic fashion [19]. It has also seen in a petite strain of *S. cerevisiae*, involving the selective activation of cryptic Pol II promoters from episomal rDNA elements [20]. The rDNA tandem array, concentrated in nucleoli of yeast where Pol I is active, is well established as a gene silencing region for Pol II activity [21]. It differs from mating loci and telomere silencing regions, in that active suppression of Pol II coexists with highly active transcription by Pol I. While a number of mechanisms have been proposed for this paradoxical observation, multiple observations, combined with reporter *mURA3* gene integration studies have led to a model of “reciprocal silencing” [22]. That is, chromatin conditions favoring Pol I, decrease or silence Pol II and vice versa. The Pol I transcribed rDNA repeats are separated by non-transcribed sequences (NTS) separated by the 5S rRNA gene. Molecular studies have localized rRNA transcription silencing of Pol II to these interweaving sequences. This is where NAD^+^-dependent histone deacetylase Sir2, as part of the RENT complex is attracted and concentrated, leading to repressive chromatin structure changes [23]. However, still Pol II can gain access even to the non-transcribed sequences as indicated by its ability to copy non-coding RNAs [24[. Our data point to another example of Pol II escaping silencing and participating in rRNA production for the cell. Target of rapamycin (TOR) signal transduction pathway regulates ribosome production including the transcription and processing of 35S rRNA [25]. As nutritional sources of the cell ebb or when exposed to rapamycin, inhibition of TOR develops, decreasing Pol I activity and eventually displacing it from the nucleolus [26]. At some point Pol II appears to get access by a hitherto unknown mechanism to rDNA, downstream from the Sir2 silenced NTS. Northern blotting of these molecules show them to be similar in size to processed ones and oligo ligation and amplification confirms that it is indeed at the processing site for both for 18S and 25S. This raises the possibility that like the processome, [27] once this access develops, small nucleolar RNAs with complementarity at or near the start sites guide Pol II to the sites and function as primers for the enzyme for further downstream transcription. This capacity to produce 18S and 25S another way, appears to function as a back-up system for the cell during unfavorable nutritional states to maintain some capacity for protein production. Our finding previously that such molecules were incorporated into ribosomes^3^ further supports this idea. Indeed *C. albicans* expresses genes specifically in the stationary phase that play important roles in pathogenesis [28]. Based on our new findings, the polyadenylated 18S and 25S molecules we have seen previously in *C*. *albicans* [9, 10] are the same molecules we are reporting now. It is of interest that we have found similar exonuclease resistant 18S and 25S molecules in *S. cerevisiae* and the fission yeast *Schizosaccharomyces pombe* [3], and polyadenylated 18S and 25S rRNA molecules have been reported for both [29, 30]. Furthermore, finding of polyadenylated rRNAs in several *Leishmania* species [31] indicates that our findings related to Pol II being involved in rRNA synthesis may be more widespread at least in monocellular eukaryotes.

## Methods

### Organisms

*Candida albican*s SC5314 (purchased from ATCC MYA 2876) was maintained in 50% glycerol in YPD broth (2% w/V tryptone, 1% w/v yeast extract, 2% w/v dextrose) at −80°C. Cells were activated in YPD broth at 30°C and maintained on Sabouraud dextrose agar at 4°C, passaged every 4–6 weeks up to 4–5 times. Yeasts were lifted from agar surface and grown in YPD broth for variable length of times at 30°C. Yeast cell concentrations were established by counting with a hemocytometer.

### RNA Isolation

Cells were collected by centrifugation, washed with sterile phosphate buffered saline (PBS) and were put on ice pending total RNA extraction. Cells were disrupted with RNase-free zirconia beads and RNA was isolated using Ambion RiboPure RNA Purification kit for yeast (Ambion/ThermoFisher) according to the manufacturer’s instructions. RNA quantification was done using a Qubit 2.0 fluorometer.

### Immunoblotting

RNA was separated on pre-fabricated formaldehyde agarose gels (Lonza) and stained with SYBR Gold Nucleic Acid Gel Stain (Life Technologies) for 30 minutes. Gel images were captured with a digital camera (Canon Vixia HFS30). Immuno-Northern blotting [32] was performed with some modifications. RNA was transferred by electro-blotting (Thermo Scientific Owl Hep-1) to a positively charged nylon membrane (Life Technologies) in 0.5 × TBE (standard Tris/Borate/EDTA buffer). The RNA was cross-linked to the membrane using UV (Stratagene UV Crosslinker). Membrane was blocked with 10% Block Ace™ (Bio-Rad) for 30 minutes at 25°C, followed by the addition of anti-7-methylguanosine (m7G) monoclonal antibody (MBL), diluted 1:1000 in 10% Block ACE™ (Bio-Rad) and incubated for 24 hours at 4° C. Goat anti-mouse conjugated to HRP was added to the membrane at 1:5000 in blocking solution for 30 minutes at 25°C. The Supersignal ™ West Femto (Thermo Scientific) chemiluminescence substrate was used to detect the HRP signal. Film was developed with the SRX-101A Konica film processor.

### Cap Binding Protein Assay

One μg of total RNA from *C. albicans* was incubated with 2 μg of recombinant human EIF4E protein fused to His-tag at N-terminus (Creative BioMArt) in binding buffer (25mM Tris, pH 8.0, 150mM NaCl, 1mM DTT, 5mM imidazole) and incubated at 4°C overnight. HisPur™ Ni-NTA magnetic beads (ThermoFisher Scientific) were added to the RNA-EIF4E mixture and placed on ice for 10 minutes. A magnetic stand was used to collect the beads after three washes with binding buffer. Capped RNA was eluted with 200 mM imidazole buffer. Finally, a phenol chloroform extraction was done in order to remove EIF4E protein off the eluate.

### 5’-End of rRNA Exonuclease Resistance

Total RNA was treated with a processive 5′→3′ exonuclease (Terminator, Epicentre) following the manufacturer’s protocol using Buffer B. The ratio of enzyme to substrate used was 1 U per 1 μg of RNA to ensure adequate cleavage.

### TOR inhibition Assay

To actively inhibit TOR, rapamycin (Sigma Aldrich) was added to fresh YPD to a final concentration of 1ug/ml. *C. albicans* were incubated in this solution at concentration of 1×10^6^ cells/ml for 60 minutes at 30°C with constant shaking. After incubation cells were washed with PBS and used for the appropriate assay.

### Decapping Assays

Cap-Clip™ acid pyrophosphatase (Cellscript) was used according to manufacturer instructions to remove the 5’-terminal m^7^GpppG “cap” from RNA samples. Verification of cap removal was done by gel electrophoresis and immuno-precipitation with cap binding protein.

### RNA Immunoprecipitation

RNA immunoprecipitation (RIP) was performed on mid-log and stationary RNA with anti-m7G-Cap mAb and protein A/G magnetic beads to purify the antibody-capped RNA complex. RNA was extracted with Direct-zol™ (Zymo Research). RT-qPCR was carried out using 18S and 25S specific primers. Positive and negative controls were actin and ITS-1 respectively.

### RNA precipitation using cap binding protein

RNA precipitation was performed on mid-log and stationary RNA with his-tagged cap binding EIF4E protein and Ni-NTA magnetic beads. RNA was eluted off the beads with 200 mM imidazole solution. RT-qPCR was done using 18S (PA-PB) and 25S (PC-PD) specific primers. Positive and negative controls were actin (PG-PH) and ITS-1 (PE-PF) respectively. See S1 Table for primer sequences.

### Native Chromatin immunoprecipitation (ChIP) analysis

C. *albicans* (1×10^6^ c/mL) were grown at 30°C for 16 hours (stationary) in a 500 mL YPD. Crosslinking was done by adding formaldehyde to the culture and incubated at RT for 20 minutes with gently swirling. After that, 37.5 mL of 3M glycine, 20mM Tris was added and incubated for 5 minutes. Cells were pelleted at 2000 rpm for 5 minutes and washed twice with 200 mL cold TBS (20mM Tris-HCl, pH 7.5, 150 mM NaCl) and once with 10 mL cold FA lysis buffer (100 mM Hepes-KOH, pH 7.5, 300 mM NaCl, 2 mM EDTA, 2% Triton X-100, 0.2 % Na Deoxycholate)/0.1% SDS. Pellets were resuspended in 1 mL cold FA lysis buffer/ 0.5% SDS. Cells were broken up by Zirconia bead (Ambion) vortexing. Chromatin isolation and shearing were done following Keogh and Burtowski [33]. Isolation of protein/DNA fragments specific for RNA polymerase II were selected with the ChromaFlash High Sensitivity ChIP Kit (Epigentek) with antibody specific for RNA polymerase II c-terminal component, following the manufacturer’s instructions. PCR analysis was done to confirm the presence of protein/DNA complexes containing 18S and 25S RNA specific sequences. PCR amplicons were sequenced (Laragen Inc.) using the same reverse primers as the ones utilized for the PCR.

### Transcription start site analysis

Components of the GeneRacer™ Kit (Life Technologies) were used to obtain new RNA transcription sites. A 44-bp oligo was ligated to previously treated with CIP and Cap-Clip™ (Cellscript) RNA using T4 RNA Ligase (Ambion). After ligation, RNA was precipitated and reverse transcription was performed using ProtoScript^®^ II Reverse Transcriptase (New England Biolabs) with specific reverse primers (Supplementary Table 1). A PCR assay was done with the cDNA as template. Primers utilized for the PCR were GeneRacer™ 5’ Primer and 18S and 25 S specific reverse primers (Supplementary Table 1). PCR products were electrophoresed on a 1% agarose (Lonza) gel at 90 mV and sent for sequencing (Laragen).

### Polyadenylation Analysis

Polyadenylated total RNA or CBP precipitated RNA was isolated using a Poly(A) RNA Selection Kit (Lexogen) [34]. Briefly, 5 μg of denatured RNA were incubated with magnetic oligodT beads for 20 minutes at 25°C, followed by 3 washes using a magnetic stand to separate the beads from the solution. RNA was eluted with water and used for subsequent experiments, such as immunoblotting or reverse transcription. Precipitated RNA was reverse transcribed with poly-T primers (PI and PJ). PCR products were amplified with the same poly-T primers and a second primer (PM or PN). Amplicons were sent for sequencing using forward primers PK or PK

### Immunofluorescence Microscopy

We followed the protocol by Wollinski and Kohlwein [35]. 1x 10^6^ cells of mid-log and stationary *C. albicans* were collected and washed 3 times with RNase-free water. After final wash, the pellet was suspended in spheroplast solution (1M sorbitol, 15mM EDTA pH 8.0, 50mM DTT), followed by centrifugation at 1500 rpm and suspension in sorbitol/citrate buffer (1M sorbitol, 1mM EDTA, 10mM sodium citrate buffer pH 5.8). Zymolase (Sigma) was then added at 300 U/ml and cells were incubated for 10 minutes (mid-log cells) and 60 minutes (stationary cells) at 30°C. Spheroplasts were gently washed with PBS and applied on a poly-L-lysine (Sigma) coated slide. After 10 minutes, cold methanol was added onto the slide and rapidly washed with cold acetone. Slides were dried at room temperature and immediately PBS + 1% BSA was added to rehydrate the cells. Mouse monoclonal antibodies raised against polymerase II (Epigentek) and Nop1p (Santa Cruz Biotechnology) at a 1:1000 dilution in PBS + 1% BSSA were incubated with the slide overnight at 4°C. Slide was gently washed 3 times with PBS and goat-anti-mouse antibodies conjugated to Alexa 488 and Alexa 568 (Invitrogen) were added at a 1:1000 dilution onto the slide and incubated for 60 minutes at 25°C. Slide was washed 3 times with PBS followed by addition of 20μM DRAQ5™ fluorescent probe (Thermo Scientific).

Imaging of immune-stained cell samples was performed on a Olympus FV1000D laser confocal microscope (Olympus Life Science Solutions, Center Valley, Pennsylvania, USA) in combination with Olympus Software Fv10-ASW 0.4 imaging software.

## Reference

1. Woolford Jr. JL, Baserga SJ. Ribosome biogenesis in the yeast Saccharomyces cerevisiae. Genetics. 2013;195: 643–681.

2. French SL, Osheim YN, Cioci F, Nomura M, Beyer AL. In exponentially growing Saccharomyces cerevisiae cells, rRNA synthesis is determined by the summed RNA polymerase I loading rate rather than by the number of active genes. Mol Cell Biol. 2003;23: 1558–1568.

3. Fleischmann J, Rocha MA. Nutrient depletion and TOR inhibition induce 18S and 25S ribosomal RNAs resistant to a 5’-phosphate-dependent exonuclease in Candida albicans and other yeasts. BMC Mol. Biol. 2018:; doi:10.1186/s12867-018-0102-y.

4. Kullberg BJ, Arendrup MC. Invasive Candidiasis. N Engl J Med 2015; 373: 1445–1456.

5. Warner JR. Synthesis of ribosomes in Saccharomyces cerevisiae. Microbiol Rev. 1989; 53: 256–271.

6. Jones T, Federspiel NA, Chibana H, Dungan J, Kalman S, Magee BB, et al. The diploid genome sequence of Candida albicans. Proc Natl Acad Sci USA. 101, 7329–7334 (2004).

7. Pendrak M, Roberts DD. Ribosomal RNA processing in Candida albicans. RNA. 2011;17: 2235–2248.

8. Adamo A, Pinney JW, Kunova A, Westhead DR, Meyer P. Heat stress enhances the accumulation of polyadenylated mitochondrial transcripts in Aradopsis thaliana. PLoS One. 2008; doi:10.1371/journal.pone.0002889.

9. Fleischmann J, Liu H, Wu CP. Polyadenylation of ribosomal RNA by Candida albicans also involves the small subunit. BMC Molecular Biology. 2004; https://bmcmolbiol.biomedcentral.com/articles/10.1186/1471-2199-5-17.

10. Fleischmann, J, Liu H. Polyadenylation of ribosomal RNA by Candida albicans. Gene. 2001; 265: 71–76.

11. Hirose Y, Manley JL. RNA polymerase II is an essential mRNA polyadenylation factor. Nature. 1998;395: 93–6.

12. Noe-Gonzalez M, Sato S, Tomomori-Sato C, Conaway JW, Conaway RC. CTD-dependent and - independent mechanisms govern co-trancriptional capping of Pol II transcripts. Nat Commun. 2018; doi:10.1038/s41467-018-05923-w.

13. Sukarieh R, Sonenberg N, Pelletier J. Nuclear assortment of eIF4E coincides with shut-off of host protein synthesis upon poliovirus infection. Journal of General Virology. 2010;91: 1224–1228.

14. Wooden J, Ciborowski P. Chromatin Immunoprecipitation for Human Monocyte Derived Macrophages Methods. 2014;70: 89–96.

15. Sirri V, Urcuqui-inchima S, Roussel P, Hernandez-Verdun D. Nucleolus: the fascinating nuclear body. Histochem Cell Biol. 2008;129: 13–31.

16. Russell J, Zomerdijk J. CBM. The RNA polymerase I transcription machinery. Biochemical Society Symposium. 2006;73: 203–16.

17. Pringle JR, Adams AEM, Drubin DG, Haarer BK. Immunofluorescence methods for yeast. 1991;194: 565–602.

18. Tollervey D, Lehtonen H, Carmo-Fonseca M, Hurt EC. The small nucleolar RNP protein NOP1 (fibrillarin) is required for pre-rRNA processing in yeast. EMBO J. 1991;10: 573–83.

19. Vu L, Siddiqi I, Lee B, Josaitis CA, Nomura M. RNA polymerase switch in transcription of yeast rDNA: role of transcription factor UAF (upstream activation factor) in silencing rDNA transcription by RNA polymerase II. Proc Natl Acad Sci USA. 1999;96: 4390–4395.

20. Conrad-Webb H, Butow RA. A polymerase switch in the synthesis of rRNA in Saccharomyces cerevisiae. Mol Cell Biol. 1995; 15: 2420–8.

21. Gartenberg MR, Smith JS. The Nuts and Bolts of Transcriptionally Silent Chromatin in Saccharomyces cerevisiae. Genetics. 2016; 203: 1563–1599.

22. Cioci F, Vu L, Eliason K, Oakes M, Siddigi IN, Nomura M. Silencing in yeast rDNA chromatin: reciprocal relationship in gene expression between RNA polymerase I and II. Molecular Cell. 2003;12: 135–145.

23. Huang J, Moazed D. Association of the RENT complex with nontranscribed and coding regions of rDNA and a regional requirement for the replication fork block protein Fob1 in rDNA silencing. Genes Dev. 2003;17: 2162–2176.

24. Li C, Mueller JE, Bryk M. Sir2 represses endogenous polymerase II transcription units in the ribosomal DNA nontranscribed spacer. Mol. Biol. Cell. 2006; 17: 3848–3859.

25. Mayer C, Grummt I. Ribosome biogenesis and cell growth: mTOR coordinates transcription by all three classes of nuclear RNA polymerases. Oncogene. 2006; 25: 6384–6391.

26. Tsang CK, Bertram PG, Ai W, Drenan R, Zheng SXF. Chromatin-mediated regulation of nucleolar structure and RNA Pol I localization by TOR. The EMBO Journal. 2003;22: 6045–6056.

27. Henras AK, Plisson-Chastang C, O’Donohue MF, Chakraborty A, Gleizes PE. WIREs RNA. 2015;6: 225–242.

28. Uppuluri P, Chaffin WL. Defining Candida albicans stationary phase by cellular and DNA replication, gene expression and regulation Molecular Microbiology.2007; 64: 1572–1586.

29. Kuai L, Fang FJ, Butler S, Sherman F. Polyadenylation of rRNA in Saccharomyces cerevisiae. Proc Natl Acad Sci USA. 2004; 101: 8581–8586.

30. Win TZ. Draper S, Read RL, Pearce J, Norbury CJ, Wang SW. Requirement of Fission Yeast Cid14 in Polyadenylation of rRNAs. Mol Cell Biol. 2006;26: 1710–1721.

31. Decuypere S. Vandesompele J, Yardley V, De Donckeri S, Laurent T, Rijal S, et al. Differential polyadenylation of ribosomal RNA during post-transcriptional processing in Leishmania. Parasitology. 2005;131: 321–9.

32. Mishima E, Jinno D, Akiyama Y, Itoh K, Nankumo S, Shima H, et al. Immuno-Northern blotting: detection of RNA modifications by using antibodies against modified nucleosides. PLoS ONE. 2015 https://doi.org/10.1371/journal.pone.0143756.

33. Keogh MC, Buratowski S. Using chromatin immunoprecipitation to map cotrascriptional mRNA processing in Sacharomyces cerevisiae. Methods Mol Bio. 2012;257: 1–6.

34. Salles FJ, Richards WG, Strickland S. Assaying the polyadenylation state of mRNAs. Methods. 1999;17: 38–45.

35. Wollinski H, Kohlwein SD. Single yeast cell imaging. Methods Mol Biol. 2014;1205:91–109.

