## Supplemental material for "RNA Polymerase II is involved in 18S and 25S ribosomal RNA transcription, in *Candida albicans*"

**S1 Fig: Dot blot analysis showing highest affinity for m7G-capped RNA. Obtained from MBL (www,mblintl.com).**

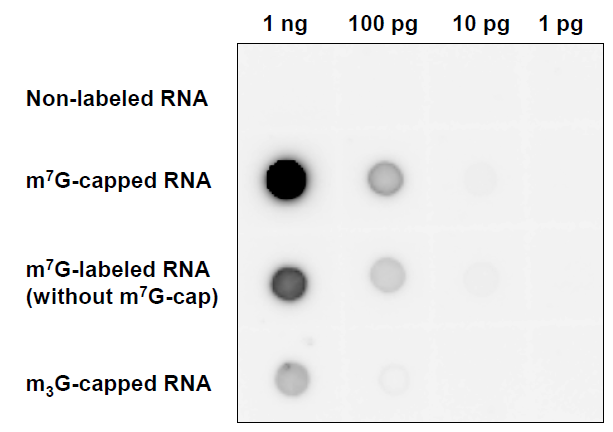

**S2 Fig: Dot blot analysis showing m7Gppp blocking m7G-cap-mAb**
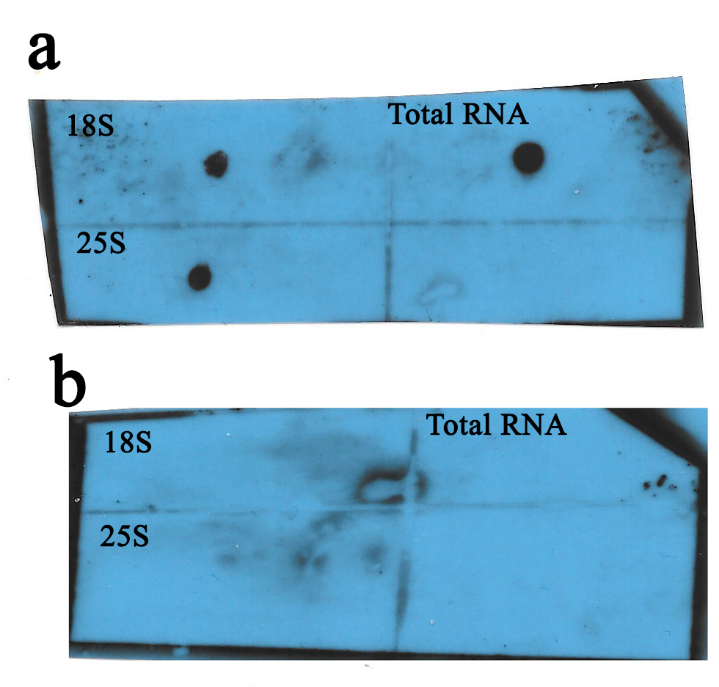

(A, B) 18S and 25S gel purified from RNA along with total RNA were UV cross linked to nylon membrane. (A) blot was detected with m7G-cap mAb. (B) blot was detected with m7G-cap mAb previously incubated with m7Gpp (Sigma Aldrich).

**S1 Table. List of primers used in all the experiments.**

| Primer designation | Primer Name | Sequence (5’-3’) | Experiment |
| --- | --- | --- | --- |
| PA | 18SfwdqPCR | AACGGCTACCACATCCAAGG | RT-qPCR, ChIP |
| PB | 18SrevqPCR | CACCAGACTTGCCCTCCAAT | RT-qPCR, ChIP, start sites |
| PC | 25SfwdqPCR | CAGGGATTGCCTCAGTAGCG | RT-qPCR, ChIP |
| PD | 25srevqPCR | CCCTCTGTGACGTCCTGTTC | RT-qPCR, ChIP, start sites |
| PE | ITS-1 fwd | AGCTGATTTGCTTAATTGCACCAC | RT-qPCR |
| PF | ITS-1 Rev | GACTATTAGTAATAATCTGGTGTGAC | RT-qPCR |
| PG | Actin fwd | GCTCCAAGAGCTGTTTTCCCA | RT-qPCR |
| PH | Actin Rev | TTTGGATTGGGCTTCATCACC | RT-qPCR |
| PI | DT12mr | TCGGCGAGCTCCGCGGCTTTTTTTTTTTT | PolyA RT |
| PJ | DT30mr | CACTCGAGCGCGCGGTTTTTTTTTTTTTTTTTTTTTTTTTTTTTT | PolyA RT |
| PK | 18sFwd(b) | CTGGGGATAGAGCATTGTAATTGTT | PolyA PCR, ChIP |
| PL | 25S-3’ Fwd | GCAGTCAAGCGTTCATAGCG | PolyA PCR, ChIP |
| PM | P3mr-25S | GTCGCTGGACCATAGCAGGCTGGCAACG | PolyA PCR |
| PN | P1mr-18S | TCGATGGAAGTTTGAGGCAATAACAGGTCTGTG | PolyA PCR |
| PO | 18SFwd(a) | GCCAGTAGTCATATGCTTGTC | ChIP |
| PP | 18SREV_9 | ACCGATCCCTAGTCGGCATA | ChIP |
| PQ | 18S-3’_Rev_end | ATCCTTCCGCAGGTTCA | ChIP |
| PR | 25SFwd(a) | ATCAGGTAGGACTACCCGCTG | ChIP |
| PS | 25SREV_2 | GCCATAAGACCCCATCTCCG | ChIP |
| PT | 25S-3’_Rev_end | AATCAGACAACAAGAGCTTAA | ChIP |
| PU | GeneRacer Oligo | CGACUGGAGCACGAGGACACUGACAUGGACUGAAGGAGUAGAAA | Start sites |
| PV | GeneRacer 5’ | CGACTGGAGCACGAGGACACTGA | Start sites |
| PW | 25S-5’_Rev_Start | TCCAAACCGATGCTGG | Start sites |
| PX | 18S-5’_Rev_Start | AGCATGTATTAGCTCTAGAATTACC | Start sites |
| PY | 25S_rev_short | GGCAATCCCTGTTGGTTTCTT | Start sites |
| PZ | 18S_rev_short | CGCAGTTTCACTGTATAAATTGCTTATACTT | Start sites |
